# Confirmation Bias Exists in the Face of False Information

**DOI:** 10.64898/2026.05.07.723487

**Authors:** Hamid Razi, Thomas Sambrook, Neil Garrett

## Abstract

Confirmation bias impacts judgments and decisions across a range of domains including finance, policy and science. Here we examine whether explicitly labelling information as true or false disrupts a core underlying computational mechanism that can generate this pervasive bias - asymmetric learning. Human participants (Study 1: N=47; Study 2: N=57) completed a 2 alternative forced choice (2AFC) task previously used to test for the presence of confirmation bias. Participants made choices between pairs of options that could win or lose money and received either factual or counterfactual feedback after each choice. We introduced a key novel feature into the task - providing explicit cues that signalled to participants whether feedback they had seen was true (verified) or false (debunked). Learning in response to feedback was attenuated under false compared to true labels but was present under both. Fitting participants choices to computational models enabled us to examine how sensitivity to the feedback varied as a function of both the label (true/false) and confirmation (confirmatory/disconfirmatory). This revealed a distinct pattern of learning rates typical of confirmation bias (enhanced learning from positive prediction errors for chosen options and from negative prediction errors for unchosen options) in response to both true and false labels. The findings highlight how confirmation bias plays an important role in the effectiveness of interventions designed to verify true and/or debunk false claims. Verification is less likely to succeed when information disconfirms prior beliefs. Conversely, debunking false claims is unlikely to succeed when the information confirms one’s prior beliefs.

## Introduction

When a financial trader downplays a price increase of shares they recently sold all the while boosting their ego from a similar increase in shares they recently bought, they are applying a well-known learning bias prevalent in decision making. Across a wide range of domains, information consistent with past choices and judgments is integrated into beliefs to a greater extent than information that challenges them. This phenomenon, known as *confirmation bias* (Bronfman et al., 2015; Klayman & Ha, 1987; Nickerson, 1998; Talluri et al., 2018), impacts a range of domains, ranging from finance (Park et al., 2010) to science (Cheng, 2018; Darley & Gross, 1983) to politics (McClung Lee, 1949).

A computational mechanism that has been shown to give rise to confirmation bias is the differential use of prediction errors. Prediction errors formally capture how surprising a new piece of information is as the degree to which the information deviates from prior expectations (Sutton & Barto, 2018). In formal learning models, beliefs change as a proportion of this surprise signal; but the degree to which this happens is determined by a learning rate. It has been shown that allowing learning rates to differ according to whether new information confirms versus disconfirms past decisions gives rise to confirmation bias (Lefebvre et al., 2022; Palminteri, Lefebvre, et al., 2017) as information that confirms past choices is amplified, whilst information that undermines them is ignored. This mechanism – a form of *asymmetric learning* - goes against classic normative theories from economics (Neumann & Morgenstern, 1944), machine learning (Russell & Norvig, 2020), and psychology (Körding & Wolpert, 2004; Maslow, 1950). However, a raft of neurobiological (Dabney et al., 2020; Garrett et al., 2014; Lefebvre et al., 2017; Sharot et al., 2011) and computational (Garrett & Daw, 2020; Lefebvre et al., 2017; Palminteri, Lefebvre, et al., 2017) evidence converge to suggest that the process of updating beliefs in the face of new information does indeed involve prediction errors changing beliefs to differing degrees, depending on whether they signal one has made the right or the wrong decision with some evidence suggesting that this strategy that can actually be optimal in certain contexts (Cazé & van der Meer, 2013; Lefebvre et al., 2022; Schubert et al., 2024).

Whilst the computational principles that give rise to confirmation bias have been established, much of the theory and empirical groundwork has been confined to cases in which information is accurate but no explicit cues are provided regarding its reliability (Chierchia et al., 2023; Lefebvre et al., 2022; Palminteri, Lefebvre, et al., 2017; Palminteri, 2023; Rollwage et al., 2020; Rollwage & Fleming, 2021; but see Vidal-Perez et al., 2025). In these paradigms, learners implicitly assume the veracity of feedback, and as such, existing accounts of confirmation bias have largely been developed in environments where the accuracy of information is taken for granted rather than explicitly interrogated. In contrast, much of our everyday experience involves gathering and processing information which is later verified or falsified. Understanding how people learn when the reliability of information is explicitly resolved raises the question of whether confirmation bias persists under these conditions and whether it arises from similar computational principles. This question is increasingly pressing in modern information environments in which digital platforms prioritise engagement over accuracy, leading to a proliferation of inaccurate content online (Lewandowsky et al., 2017) which tends towards reinforcing rather than challenging pre-existing beliefs and views (Del Vicario et al., 2016; Jiang et al., 2021).

To address this, we adapted a novel learning paradigm that extends the two-alternative forced choice (2AFC) task previously used to test for choice confirmation bias (Lefebvre et al., 2022; Palminteri, Lefebvre, et al., 2017) by making feedback reliability explicit. In the task (**Figure 1(a)**), participants (Study 1: N=47; Study 2, replication: N=57) made choices between pairs of options (abstract symbols) and were then shown an outcome (gain/loss) for one of the two options (either the option chosen or the unchosen option). Cues then indicated whether the outcome revealed had been genuine (verified) or false (debunked) allowing us to directly examine how learning unfolds in the face of explicitly reliable versus unreliable information. We used this setup to examine learning in the context of both loss avoidance and gain maximisation in order to disassociate effects driven solely by outcome valence. Crucially, participants were provided with either counterfactual (outcome shown for the unchosen option) or factual (outcome shown for the chosen option) feedback after each choice. This feature is critical in order to be able to disassociate confirmation bias - preferential updating from choice-consistent evidence - from positivity bias – a general tendency to overweight positive compared to negative prediction errors independent of choice (Palminteri & Lebreton, 2022; Schubert et al., 2024). Computationally, confirmation bias predicts asymmetric updating conditional on choice (enhanced learning from positive prediction errors for chosen options and from negative prediction errors for unchosen options), whereas positivity bias predicts uniformly greater learning from positive prediction errors irrespective of choice. This distinction cannot be resolved in paradigms that provide only factual feedback (Lefebvre et al., 2017; Vidal-Perez et al., 2025).

**Figure 1.**
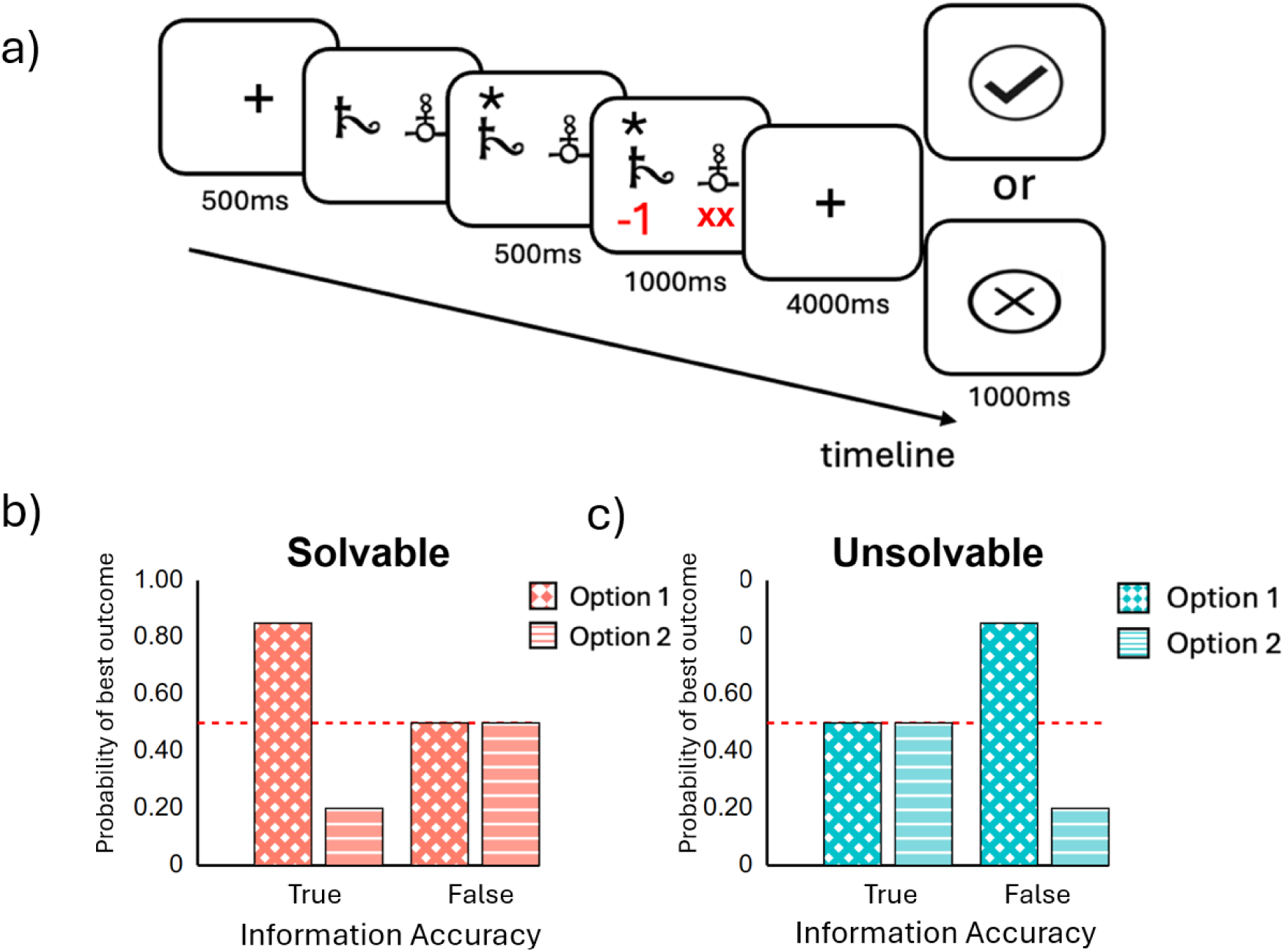
Experimental Design. **(a)** Participants made choices between four pairs of options (abstract symbols) and received an outcome (win money in the gain pairs, lose money in the loss pairs) about one of the two options (the option chosen or the unchosen option). An information accuracy cue (tick or cross) then indicated whether that outcome had been true (tick) or false (cross). **(b)** For two of the pairs, in true trials, one of the two options was truly more likely to lead to the best outcome (either winning money or avoiding losing money) and outcomes were random for each option on false trials. **(c)** For the other two pairs, this pattern was reversed, such that each option randomly led to one of two outcomes. But on false trials, one option was more likely to show the better outcome (either winning money or avoiding losing money) before the false information accuracy cue was revealed.

## Methods

### Participants

A total of 70 participants were recruited online via Prolific (Palan & Schitter, 2018) for the first Study, 23 of whom were excluded; therefore, the final sample was 47 participants (mean [standard deviation] age: 30.45 [7.2]; 28 female). A total of 91 participants were recruited from the university pool for the second Study, 34 of whom were excluded, leaving the final sample of 57 participants (mean [standard deviation] age: 20.45[3.8]. Two exclusion criteria were applied to ensure data quality (Nadler et al., 2021; Peer et al., 2022; Zorowitz et al., 2023). First, participants who incorrectly answered more than one of ten catch trials were excluded (n=1 in Study 2). Second, participants showing subpar learning performance were excluded, defined as choosing the better option less than 55% of the time in solvable conditions (n=23 in Study 1, n=33 in Study 2). Study 1 participants received £3 plus a performance-based bonus ranging from £3-£6. Study 2 participants received course credits plus a performance bonus up to £3. All participants provided informed consent prior to participation. The research protocol received approval from the University of East Anglia’s ethics committee and complied with all relevant ethical guidelines.

### Behavioural task

Participants completed a novel instrumental learning task (**Figure 1(a)**). On each trial, they chose between two options and were then shown the outcome for one of the options (either the option chosen or unchosen). After this, they were shown an “information accuracy cue” which signalled whether the outcome revealed had been true (tick symbol displayed) or false (cross symbol displayed). Participants were instructed that when they were shown the false information accuracy cue, a "glitch" had occurred causing an outcome from an unrelated game to be shown, and that they should ignore this outcome as it was unrelated to the actual outcome of the current trial.

The experiment consisted of 160 trials (4 choice pairs, 40 trials per choice pair) organised into two gain and two loss contexts. Trials with reaction times below 100ms or above 4 seconds were removed from all analysis (5% in Study 1 and 7% in Study 2). In gain trials, participants chose between pairs that could receive +1 or +10 points. Loss trials involved choosing between pairs that could lose -1 or -10 points. Participants’ goal was to identify through trial and error which options more frequently provided the higher reward and less frequently incurred the highest loss. The total number of points participants had at the end of the experiment was converted into a bonus payment. 50% of trials presented true information accuracy cues (20 per choice pair) and the remainder presented false information accuracy cues. Option pairs were presented in an interleaved random order with randomised left/right positioning. As an attention check and to encourage ongoing monitoring of the information cues, ten catch trials were shown to participants at random points during the experiment. This probed participants to recall whether the previous trial had revealed a true or false information accuracy cue.

The underlying reward structure of each choice pair could take one of two forms. Two of the pairs (one gain pair and one loss pair) were ‘solvable’ (**Figure 1(b)**). For these choice pairs, one option (which we refer to as O1) had a greater chance of receiving the more favourable outcome of the two (+10 or -1). Participants could learn which was the better option therefore by attending to the true feedback. Of these pairings, false feedback was set such that both options (O1 and O2) were equally likely to lead to favourable or unfavourable outcomes before the false information accuracy cue was revealed. The remaining two choice pairs (again one gain and one loss pair) were ‘unsolvable’ (**Figure 1(c)**). For these choice pairs, both options had a 50/50 chance of receiving the more favourable or unfavourable outcome and this was reflected in true feedback. But for these pairings, false feedback was set such that one option (which we again refer to as O1) was more likely to lead to a favourable outcome before the false information accuracy cue was revealed.

The task was programmed in JavaScript using the toolbox jspych version 6.3 (Leeuw et al., 2023).

### Behavioural Analysis

To assess the degree to which participants used true and false information, we calculated the proportion of times each participant selected the truly better option of each pair (in the case of solvable contexts) or the falsely better option of each pair (in the case of unsolvable contexts). We then calculated average choice rates for each participant for Solvable and Unsolvable contexts (i.e. averaging over gain and loss pairs) and conducted one-sample t-tests against a test value of 0.5 to infer whether choice differed from chance in each context.

### Computational Models

We fitted several models that were variations of the standard Rescorla-Wagner (RW) model (Rescorla & Wagner, 1972). On each trial (t), either the value of the chosen (Qc) or unchosen (Qu) option is updated depending on which outcome is shown:

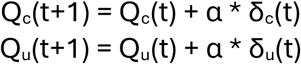

In which δ is the prediction error (δ), defined as the difference between the expected outcome and observed outcome:

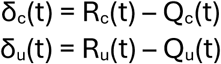

α is a learning rate determining the extent to which δ is used to update the relevant Q value. The probability of picking option A over option B at the time of choice on each trial is calculated by inputting the current Q value estimates into a standard Logit Choice rule (Softmax function):

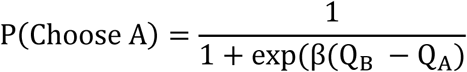

β is an inverse temperature parameter that controls the degree of stochasticity in choices (larger values of β yield more deterministic choices in favour of the option with the higher Q value, smaller values reflect more exploratory behaviour).

The contribution of each trial to the likelihood was given by the log probability of the observed choice. For a choice between the chosen (c) and unchosen (u) options:

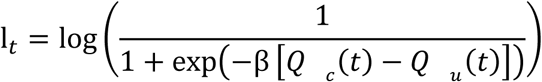

The log-likelihood for a subject was then:

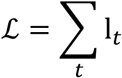

The model fitting procedure was set to minimise the negative log-likelihood, during estimation.

We fitted four models which varied in the number of learning rates (2, 3 or 4) which enabled learning to vary to differing degrees according to whether prediction errors were confirmatory/disconfirmatory and whether the feedback transpired to be true/false.

Confirmatory feedback are cases in which δ_c_ > 0 and δ_u_ < 0. Disconfirmatory feedback are cases in which δ_c_ < 0 and δ_u_ > 0. The intuition here is that both a positive prediction error generated from the option chosen and a negative prediction error generated from the option not chosen both signal one has made the correct choice (i.e. serve as instances of confirmatory feedback). Conversely, both a negative prediction error generated from the option chosen and a positive prediction error generated from the option not chosen signal one may have made an incorrect choice (i.e. serve as instances of disconfirmatory feedback).

The update rule for the four models was formulated as follows:

#### Model 1 (M1)

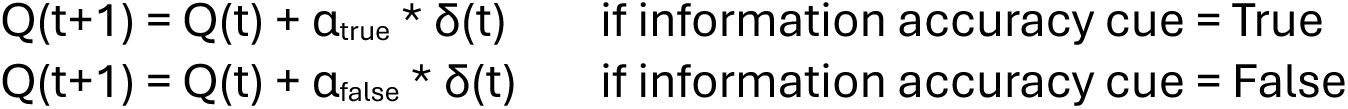

Free parameters (n=3): α_true,_ α_false,_ β

#### Model 2 (M2)

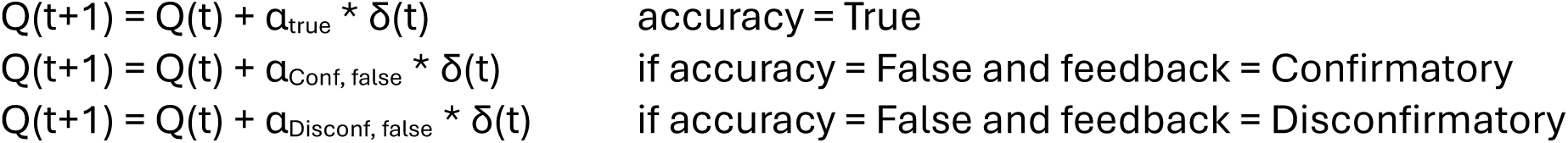

Free parameters (n=4): α_true_, α_Conf, false_, α_Disconf, false_, β

#### Model 3 (M3)

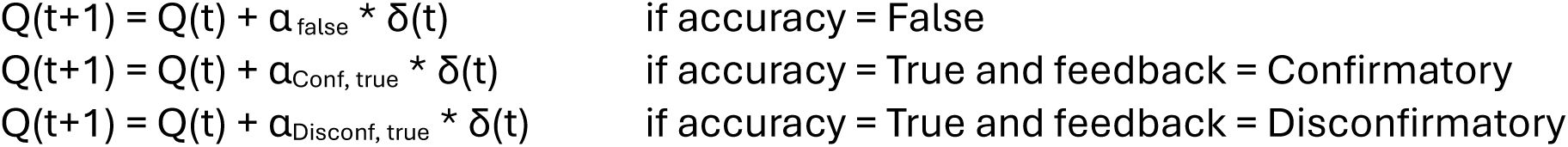

Free parameters (n=4): α_false_, α_Conf, true_, α_Disconf, true_, β

#### Model 4 (M4)

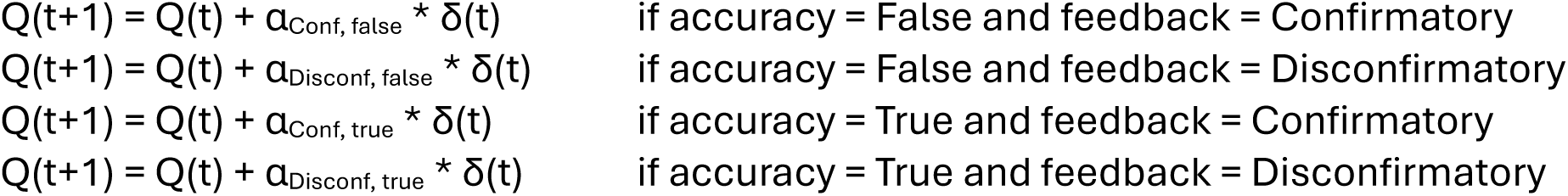

Free parameters (n=5): α_Conf, true_, α_Disconf, true,_ α_Conf, false_, α_Disconf, false_, β

We also fitted three additional models which were variations of M4 (see **Supplementary Materials** for full details). In the first, we incorporated separate learning rates for factual and counterfactual outcomes; in the second, we incorporated separate learning rates for gain and loss contexts; and, in the third, we added gradual perseveration parameters (Katahira, 2018; Sugawara & Katahira, 2021; Palminteri, 2023).

### Model Fitting Procedure

Models were fitted hierarchically by maximising the likelihood of observed choices using an Expected Maximisation (EM) algorithm (Huys et al., 2011) in Julia (v1.9.4) (Bezanson et al., 2012). In this approach, instead of fitting in isolation, the model assumes that while each participant has a unique set of parameters, these are drawn from a common group-level distribution - a Gaussian distribution defined by a group mean and variance. By doing so, the model "borrows statistical strength" from the entire group to inform each individual’s parameter estimate, down-weighting the influence of unreliable participants (Efron & Morris, 1977).

### Model Comparison

To compare model performance, we calculated subject-level leave-one-out cross-validation (LOOcv) scores. This is a method for estimating a model’s out-of-sample predictive accuracy (i.e. a model’s ability to generalize to new, unseen individuals). The process begins by temporarily holding out a single subject from the dataset. The hierarchical model is then re-fitted (using EM) using the data from all *other* subjects except the held-out subject which generates a set of cross-validated group-level parameters that are not influenced by the held-out subject’s data. These cross-validated group parameters are subsequently used as a Bayesian prior to compute the marginal likelihood of the held-out subject’s data, a score that quantifies how well the model, trained on the rest of the population, predicts the behaviour of a novel individual. Because each subject’s score is computed based on a model that was not trained on their own data, the resulting set of scores across the group can be treated as independent, which makes them suitable for subsequent classical statistical tests at the group level (e.g., t tests) and Bayesian model selection.

Subject-level LOOcv scores were submitted to the mbb-vb-toolbox in MATLAB for group-level Bayesian model selection (BMS) (Daunizeau et al., 2014). The toolbox implements a random-effects Variational Bayesian Approach (VBA). Unlike a fixed-effects analysis this random-effects approach uses a more plausible assumption: that different models may best describe different subjects (Stephan, et al., 2009). The VBA provides two key metrics for inference: model frequency and exceedance probability (XP). Model frequency estimates posterior probability that a given model generated the data for a randomly chosen subject from the population. The frequencies for all models under consideration sum to 1, which should be compared to the chance level (1 divided by the total number of models). This metric is useful for understanding population heterogeneity, as it may reveal that multiple competing models are prevalent, rather than one single winner. XP quantifies the belief that a given model is the most prevalent. For instance, a model could have the highest model frequency (e.g., 0.4) but still have a low XP if other models have similar frequencies (e.g., 0.35 and 0.25), reflecting uncertainty about which is truly the most common. An XP near 1, however, provides strong evidence for a single "winning" model, indicating a high degree of confidence that it is the most common data-generating process in the population (Rigoux et al., 2014). As an additional check, we compared LOOcv scores of the winning model to the other models using paired sample t-tests, correcting for multiple comparisons by adjusting the p value according to the number of models being compared - using the p.adjust function in base R (method = "fdr").

### Statistical Tests on the Learning Rates

Applying standard frequentist tests to individual parameter point-estimates (e.g., t-tests to compare two learning rates) from a hierarchical model fitting process like EM is statistically invalid. This is because the regularization (or "shrinkage") inherent in the fitting process violates the assumption of independence as each individual’s estimate is influenced by the group distribution. This shrinkage artificially reduces inter-subject variance, leading to a substantial inflation of the Type I error rate (i.e., false positives) (Piray et al., 2019). To overcome this challenge and be able to infer statistical differences between learning rates - following Piray and colleagues’ (2019) approach - we used hierarchical t-tests. This method operates on the posterior distribution of the group-level parameters (e.g., the group mean), which correctly reflects the uncertainty of the estimate at the population level. The test evaluates whether a credible interval for the group mean effect includes zero, based on the estimated mean and its hierarchical standard error. In order to be able to implement hierarchical t-tests we reparametrized the winning model to create parameters that directly quantified the magnitude of confirmation bias for true and false information (see **Supplementary Materials** for the equations and further details).

### Model Recovery

To verify that the different models we fitted to the choice data would be correctly inferred from the model comparison process, we conducted a model recovery analysis (Palminteri, Wyart, et al., 2017). This began with a simulation step whereby for each candidate model, a synthetic dataset for a simulated group of subjects was generated, with the number of subjects and trials mirroring the real experiment. The parameter values used for this simulation were the ranges observed in the actual data (drawn uniformly). This process resulted in a collection of synthetic datasets, one for each model. Next, we subjected each of these datasets to the full model comparison pipeline. Specifically, we fitted every candidate model to each synthetic dataset and fed the resulting LOOcv scores into the VBA toolbox to determine a winning model based on the exceedance probability. We repeated this entire simulation-and-comparison process for 50 iterations and recorded how often the model used to simulate the data was correctly identified as the best-fitting model (e.g., if data were simulated using model M1, M1 should have the highest XP value). Next, we aggregated the outcomes into a confusion matrix, which visualizes the proportion of iterations in which data generated from a specific model (rows) was correctly identified as best fit by that same model (columns). An ideal confusion matrix has high values on the diagonal (indicating successful recovery) and low values on the off-diagonals (indicating low confusion between models). This result validates that the models make sufficiently different predictions and that the comparison method is sensitive enough to identify the true underlying model (see **Supplementary Materials** for a schematic of this approach)..

### Parameter Recovery

We conducted parameter recovery for the winning model. We simulated data from the winning model using parameter values drawn randomly from uniform distributions spanning the empirically observed range. The standard hierarchical fitting procedure was then used to "recover" the parameters from the simulated data. Recovery success was assessed in two ways. First, the relationship between true and recovered parameter values was examined using Pearson correlations. We used a Perason value of 0.80 and above as evidence that a parameter is well-constrained by the data and can be estimated reliably (Daw, 2011). Second, we constructed a correlation matrix of the recovered parameters to visualise correlations between different parameters. High correlation between a pair of parameters would indicate an identifiability issue, suggesting that the model is unable to disentangle the unique contribution of each parameter to the behaviour (Wilson & Collins, 2019).

## Results

### Participants learn from true and false information

The underlying reward structure of each choice pairings in our learning task could take one of two forms (see **Methods** and **Figure 1**). In ‘Solvable’ parings, one option (O1) truly had a greater chance of receiving the more favourable outcome (+10 or -1) and false feedback was set such that both options (O1 and O2) were equally likely to lead to favourable or unfavourable outcomes. In ‘Unsolvable’ parings, both options had a 50/50 chance of truly receiving the more favourable outcome and false feedback was set such that one option (which we refer to as O1) was more likely to lead to a favourable outcome.

Analysing choice rates revealed that on average participants selected O1 – the truly better option for solvable choice pairs and the misleadingly better option for unsolvable choice pairs - over O2 for both solvable (Study 1: t(46) = 14.66, p < 0.01; Study 2: t(56) =12.50, p < 0.01) and unsolvable pairs (Study 1: t(46) = 2.64, p < 0.05; Study 2: t(56) = 3.47, p < 0.01), **Figure 2**. This suggests that participants integrated feedback from both true and false information. However, propensity to choose O1 over O2 was greater for solvable compared to unsolvable pairs (Study 1: t(46) = 5.42, p < 0.01; Study 2: t(56) = 5.48, p < 0.01), suggesting accuracy cues significantly modulated learning from feedback. As an additional check to verify that participants paid attention to information accuracy cues, on 6.25% of trials, participants were asked to report what the accuracy cue had been on the previous trial – participants correctly reported this 95% of the in Study 1 and 93% of the time in Study 2.

**Figure 2.**
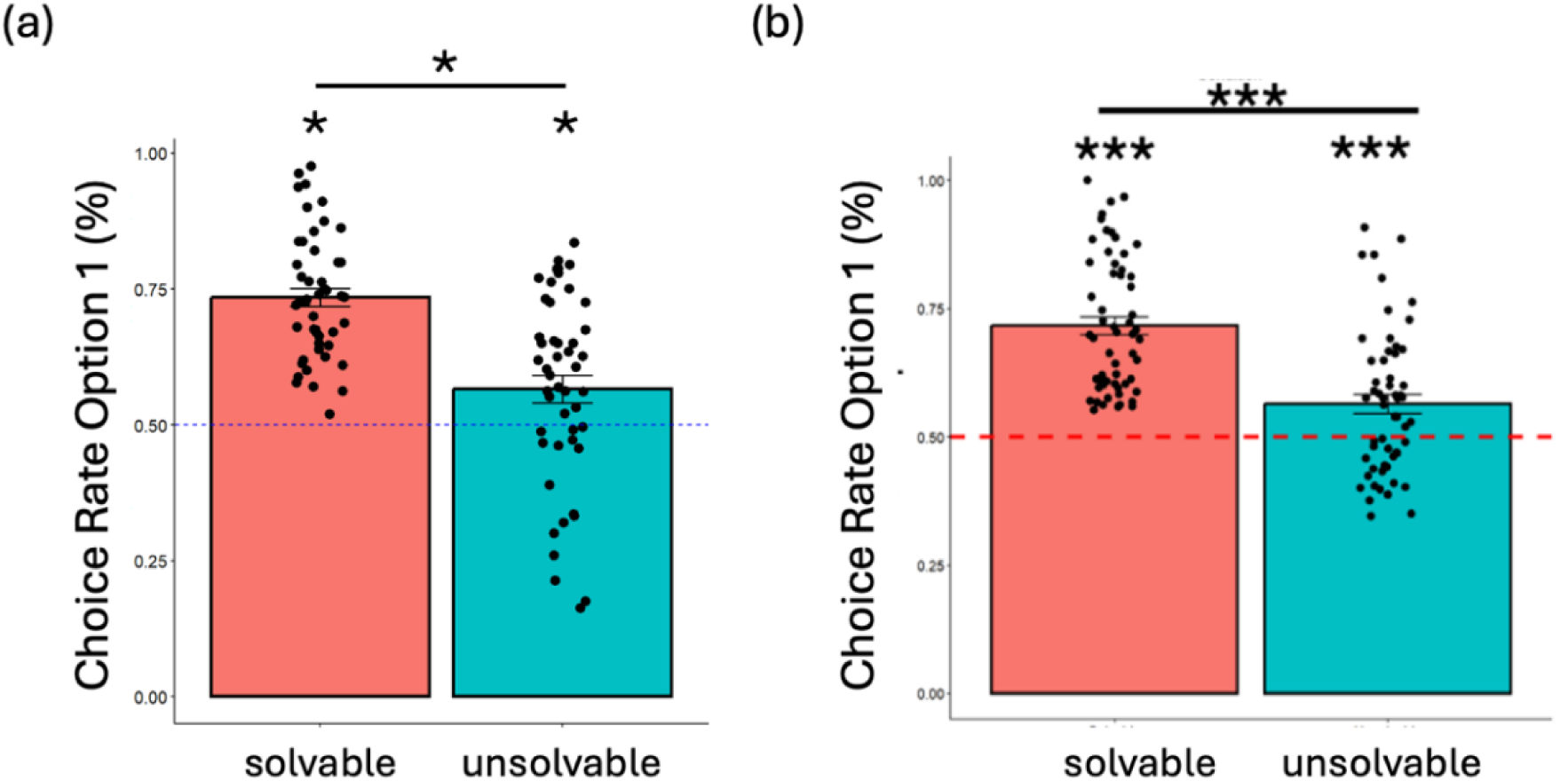
Choice Rates in (a) Study 1 and (b) Study 2. Participants opt to select option 1 - the option that provides the best outcome (+10 in the gain context or -1 in the loss context) more often relative to option 2 - to a greater degree in solvable compared to the unsolvable conditions(Study 1: t(46) = 5.42, p < 0.01; Study 2: t(56) = 5.48, p < 0.01). But, in both solvable (Study 1: t(46) = 14.66, p < 0.01; Study 2: t(56) =12.50, p < 0.01) and unsolvable (Study 1: t(46) = 2.64, p < 0.05; Study 2: t(56) = 3.47, p < 0.01) conditions, participants chose O1 to a greater degree than chance. Choice rates are averaged over gain and loss contexts. *p < 0.05, ***p < 0.001 (one-tailed test vs 0.5 or paired sample t-test as appropriate). Error bars indicate standard error of the mean (SEM).

Next, we sought to assess whether learning from true and false information exhibited confirmation bias. To do this, we fit four different computational models to the choice data (see **Methods**). These all shared the same basic structure but varied in the degree to which they parsed confirmatory versus disconfirmatory feedback and true versus false feedback. Model 1 used two learning rates: one for true and another for false information. Model 2 had three learning rates, maintaining a single rate for true information but splitting false information into separate rates for confirmatory and disconfirmatory false information. Model 3 also used three learning rates but took the opposite approach, using one rate for false information while distinguishing between confirmatory true and disconfirmatory true information. Finally, Model 4 incorporated four learning rates, providing separate rates for each combination: confirmatory true, disconfirmatory true, confirmatory false, and disconfirmatory false information.

### Four learning rate model the best fit to the data

After fitting choice data to the 4 models, we computed leave-one-out cross-validation (LOOcv) scores for each of them. Model 4 (which separated learning rates according to *both* whether feedback was true/false and whether it was confirmatory/disconfirmatory) had the lowest LOOcv score (lower scores indicate better fit) for the largest portion of participants (Study 1: 44.7%; Study 2: 42.1%) followed by Model 3 (Study 1: 25.5%; Study 2: 22.8%), and then Models 2 (Study 1: 14.9%; Study 2: 19.3%),) and 1 (Study 1: 14.9%; Study 2: 15.8%). We formally compared the LOOcv scores using a Variational Bayesian Approach (VBA) (see **Methods**). In both studies, Model 4 consistently provided the best fit to the data. Model 4 and had an exceedance probability of 1.0 in both studies and achieved the highest model frequency (∼83% in Study 1 - **Figure 3(a)** - and ∼93% in Study 2 – **Figure 3 (b)**, well above the 25% chance level) indicating strong evidence that the four-parameter learning structure of Model 4 was the best among the set of models we ran at capturing participants’ behaviour. Model Recovery verified that each of the 4 models could be reliably identified by the model fitting procedure we used (see **Methods** and **Supplemental Materials**).

**Figure 3.**
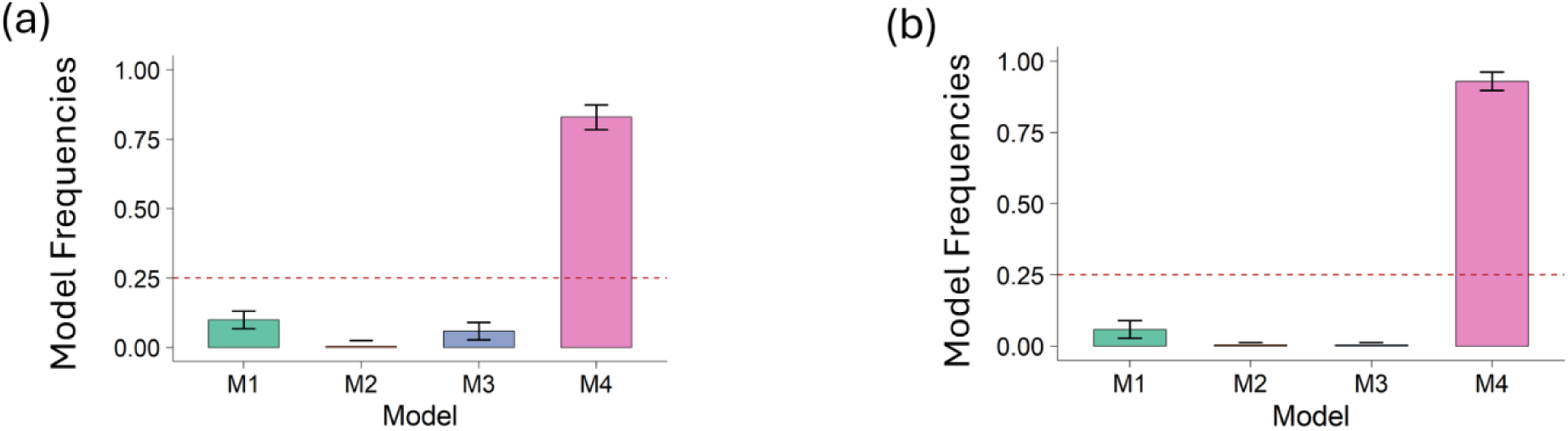
Model Comparison. **(a)** Estimated model frequencies from the VBA model comparison in Study 1. Model 4 (M4) had the highest frequency, selected for approximately 83% of participants. **(b)** Estimated model frequencies from the VBA model comparison in Study 2. Similar to Study 1, Model 4 (M4) had the highest frequency, selected for approximately 93% of participants. The model frequency reflects the proportion of the population best accounted for by each model (see **Supplementary Materials** for additional model diagnostics).

Comparing LOOcv scores using paired sample ttests (FDR corrected for multiple comparisons) revealed a similar pattern of results with Model 4 having lower LOOcv scores compared to Model 1 (t(46) = −4.75, p_adj < 0.001), Model 2 (t(46) = −3.69, p_adj <0.001), and Model 3 (t(46) = −3.07, p_adj < 0.01) in Study 1. Similarly in Study 2, Model 4 had the lowest LOOcv score compared to Model 1 (t(56) = −4.94, p_adj < 0.001), Model 2 (t(56) = −3.82, p_adj < 0.001) and Model 3 (t(56) = −3.34, p_adj < 0.01).

### Confirmation bias exists for true and false information

We then examined the pattern of learning rates from Model 4 to see if they were consistent with confirmation bias (Lefebvre et al., 2022; Palminteri, Lefebvre, et al., 2017; Palminteri & Lebreton, 2022; Schubert et al., 2024) and whether this bias was present in response to both true and false information (**Figure 4**). Examining and comparing learning rates using hierarchical t-tests (Piray et al., 2019, see **Methods**) showed that indeed participants exhibited a strong confirmation bias following false (Study 1: t(46) = 5.19, p < 0.001; Study 2: t(56) = 2.43, p = 0.01) and following true (Study 1: t(46) = 6.50, p < 0.001; Study 2: t(56) = 6.38, p < 0.001) information, learning more from confirmatory compared to disconfirmatory feedback. Comparing the bias (i.e. the difference in confirmatory versus disconfirmatory learning rates) between true and false did not show a significant difference (t(46) = 1.81, p = 0.07; t(56) = 1.11, p = 0.26). Parameter Recovery verified that each of the parameters of Model 4 could be reliably recovered by the model fitting procedure we used (see **Methods** and **Supplemental Materials**).

**Figure 4.**
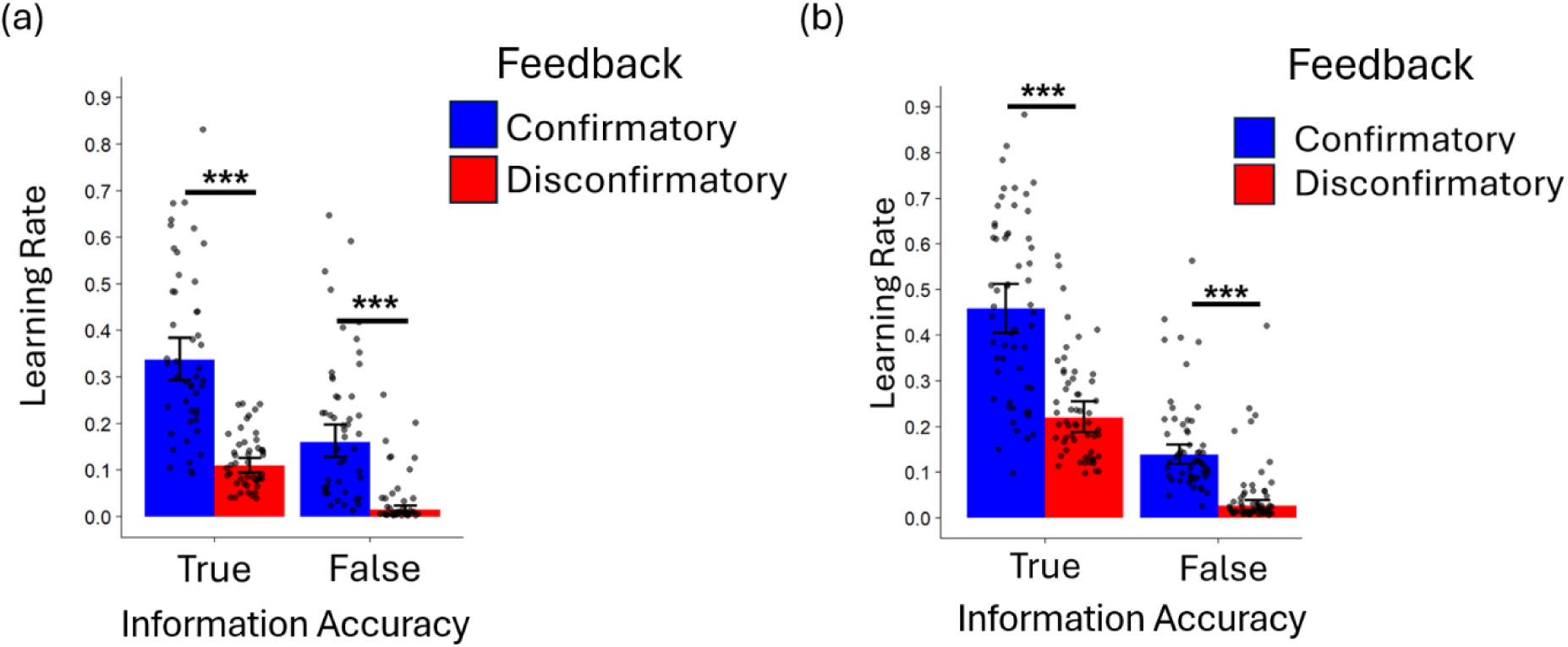
Confirmatory and Disconfirmatory Learning Rates. Learning rates from M4 (which decomposes learning into 4 types depending on the information received: Confirmatory True, Disconfirmatory True, Confirmatory False and Disconfirmatory False. (a) In Study 1, the magnitude of learning rates were higher for confirmatory versus disconfirmatory feedback (i.e. confirmation bias) both for true (t(46) = 6.50, p < 0.001, hierarchical t-test comparing α_conf_true_ with α_disconf_true_) and from false (t(46) = 5.19, p < 0.001, hierarchical t-test comparing α_conf_false_ with α_disconf_false_) information. **(b)** This pattern was replicated in Study 2 with confirmatory greater than disconfirmatory both from true (t(56) = 5.24, p < 0.001, hierarchical t-test comparing α_conf_true_ with α_disconf_true_) and from false (t(56) = 2.43, p = 0.01, hierarchical t-test comparing α_conf_false_ with α_disconf_false_) information. ***p < 0.001, *p < 0.05, hierarchical t-test. Error bars indicate hierarchical SEM.

A key feature of Model 4 (following Lefebvre et al., 2022; Palminteri, Lefebvre, et al., 2017; Palminteri & Lebreton, 2022) is that it collapses learning rates across positive prediction errors generated when feedback is received for the option chosen and negative prediction errors generated when feedback is received for the option not chosen. The rationale is that confirmation lies both in seeing one’s choice rewarded and one’s forgone choice go unrewarded (or punished). Following a similar logic, learning rates from negative prediction errors for the option chosen and positive prediction errors for the option not chosen are also combined (as both present instances of disconfirmatory evidence). More complex models that parse out learning rates for these 4 cases (positive PE chosen, negative PE unchosen, negative PE chosen and positive PE chosen) rather than collapsing them into two learning rates therefore provide a less parsimonious account of learning if sensitivity to the two confirmatory cases and sensitivity to the two disconfirmatory cases are respectively similar, as observed previously (Palminteri, Lefebvre, et al., 2017).

To verify that the underlying pattern of learning in Model 4 did indeed conform to confirmation bias we ran an additional learning model which this time separated learning out for the four different cases (positive PE chosen, negative PE chosen, positive PE unchosen and negative PE unchosen) for true and false separately (so 8 learning rates in total) and tested for an interaction between prediction error sign (positive/negative) and feedback (chosen/unchosen) separately for true and false feedback (see **Supplementary Materials** for full details of the model specification).

Consistent with confirmation bias (Palminteri, Lefebvre, et al., 2017), this revealed a significant interaction in both experiments from true (Experiment 1: t(46) = 5.95, p < ; Experiment 2: t(56) = 3.95, p < 0.001) and from false (Experiment 1: t(46) = 4.68, p < 0.001; Experiment 2: t(56) = 3.92, p < 0.001) feedback. In Experiment 1 (**Figure 5**), this was the result of higher learning rates for positive compared to negative prediction errors when receiving feedback for the option chosen. This was the case both for true feedback (t(46) = 2.59, p = 0.01) and for false feedback (t(46) = 2.20, p=0.03). But we observed the opposite pattern when participants received feedback for the option unchosen, i.e. lower learning rates for positive compared to negative prediction errors. Again, this was the case both for true feedback (t(46) = −2.75, p = 0.008) and for false feedback (t(46) = −2.47, p = 0.01). The differences were weaker in Experiment 2, but the same basic pattern replicated (see **Supplementary Materials**). This pattern of interaction we observe in both studies is the distinctive hallmark of choice confirmation bias *not* positivity bias. If it were the latter, learning would be greater for positive prediction errors compared to negative independent of chosen versus unchosen (Palminteri, Lefebvre, et al., 2017).

**Figure 5.**
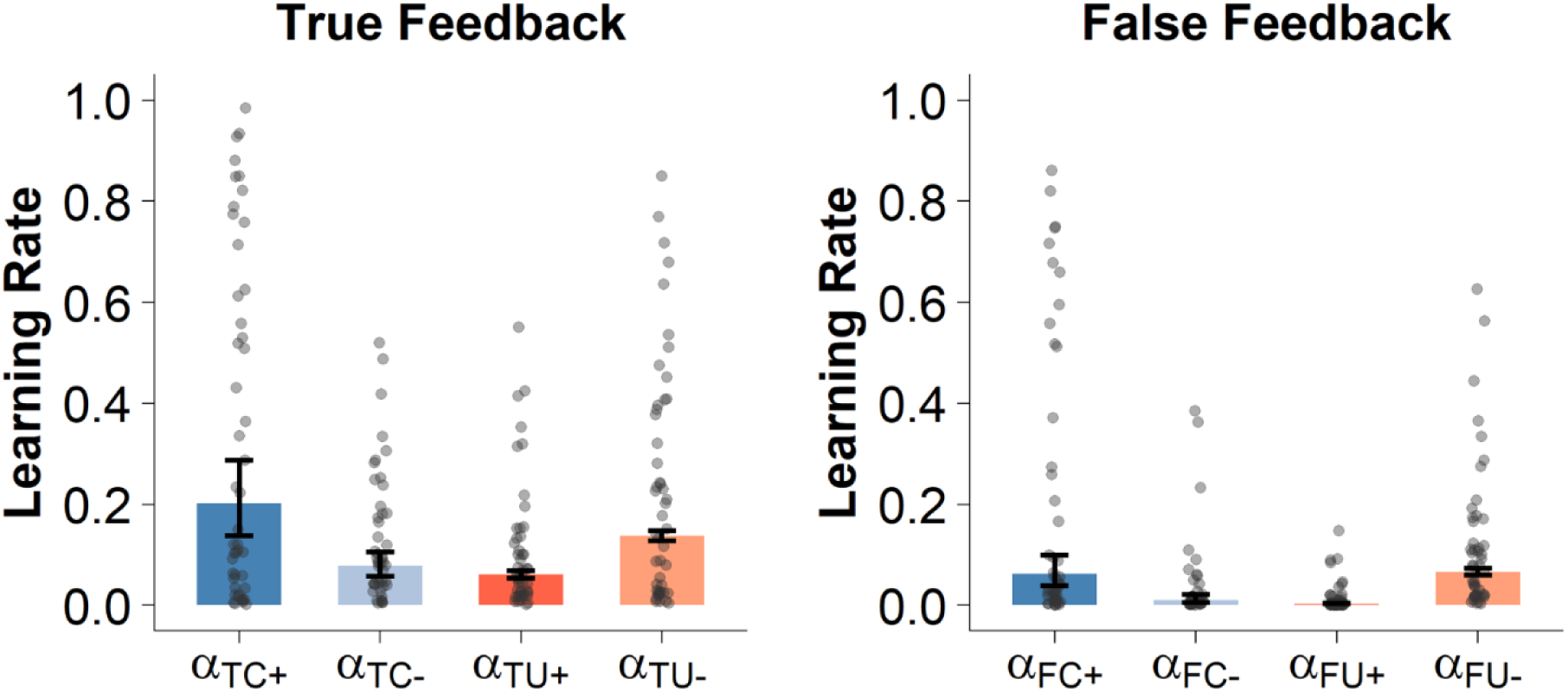
Factual and Counterfactual learning rates from positive and negative prediction errors (Experiment 1). We ran an additional model which separated learning according to whether feedback was factual (i.e. presented for the option chosen) or counterfactual (presented for the option not chosen) and the sign of the prediction error (positive or negative). There was a significant interaction which matched the pattern expected for confirmation bias in the case of (a) true (t(46) = 5.95, p < 0.001) and (b) from false (t(46) = 4.68, p < 0.001) feedback. Plotted are the learning rates from Experiment 1. See Supplementary Materials for model specification and learning rates for Experiment 2. α _TC+_ = learning rate for positive prediction errors for option chosen, true feedback; α _TC-_ = learning rate for negative prediction errors for option chosen, true feedback; α _TU+_ = learning rate for positive prediction errors for option not chosen, true feedback; α _TU-_ = learning rate for negative prediction errors for option not chosen, true feedback; α _FC+_ = learning rate for positive prediction errors for option chosen, false feedback; α _FC-_ = learning rate for negative prediction errors for option chosen, false feedback; α _FU+_ = learning rate for positive prediction errors for option not chosen, false feedback; α _FU-_ = learning rate for negative prediction errors for option not chosen, false feedback. Error bars indicate hierarchical SEM.

The key feature of learning in a confirmatory biased manner is that agents are prone to downweigh evidence that they made a mistake and should select a different option next time. It has been argued (Katahira, 2018) that - as a result – models with a single learning rate that account for choice perseverance (i.e. agents simply being more likely to repeat the previous choice independent of feedback) can account for 2AFC data just as well as models that facilitate asymmetric learning (via different learning rates). Given this, an important robustness test of confirmation bias proposed (Palminteri, 2023) is to include *both* choice perseverance and asymmetric learning in the same computational model. Accordingly, we ran a version of M4 with perseverance learning additionally controlled for (see **Supplementary Methods** for details). Comparing the learning rates from this model again revealed confirmation bias from false information both in Study 1 (t(46) = 3.13, p < 0.001) and Study 2 (t(56) = 5.24, p < 0.001). Confirmation bias from true information remained significant in Study 1 (t(46) = 2.43, p = 0.01) but not in Study 2 (t(56) = 1.15, p = 0.25).

Finally, we examined whether asymmetric learning was prevalent both in Gain (where participants learned to pick options that led to rewards) and Loss (where participants learned to pick options that avoided losses) contexts by running a version of Model 4 that separated out confirmatory and disconfirmatory learning rates for gain and loss pairs. Confirmation bias was present in Gain contexts for true (Experiment 1: t(46) = 2.51, p = 0.01; Experiment 2: t(56) = 2.46, p = 0.01) and false (Experiment 1: t(46) = 3.99, p < 0.001; Experiment 2: t(56) = 5.06, p < 0.001) information. Confirmation bias was also present in the Loss contexts both for true (Experiment 1: t(46) = 4.05, p < 0.001; Experiment 2: t(56) = 3.48, p < 0.001) and false (Experiment 1: t(46) = 4.37, p < 0.001; Experiment 2: t(56) = 6.25, p < 0.001) information (see **Supplementary Materials** for model specification and plots of the learning rates from this model in each dataset).

## Discussion

Confirmation bias - the tendency to update beliefs more readily from information that confirms rather than disconfirms prior choices – is a pervasive bias that affects decision making in a range of domains (Lord et al., 1979). Here in two separate studies, we show that this bias persists both when information is verified and when it is debunked. To show this, we designed a novel version of the classic 2AFC task in which participants received either factual or counterfactual feedback after making a choice and were then presented with an explicit cue post feedback, revealing whether the feedback had been true or false. Whilst learning attenuated following debunked (false cue) compared to verified (true cue) feedback, participants integrated feedback and adapted their decisions following both types of cues.

By including both factual and counterfactual feedback, our design allows us to distinguish confirmation bias from a simple positivity bias at the computational level. We observed greater learning from positive relative to negative prediction errors when feedback is provided for the chosen option, but the reverse pattern when feedback is provided for the unchosen option. This interaction - whereby learning tracks whether an outcome validates the correctness of one’s choice rather than its valence per se - is the hallmark of confirmation bias, and disassociates from positivity bias, which predicts greater updating from positive prediction errors irrespective of choice. The confirmation bias pattern was present for both true and false information, suggesting that the asymmetry operates over perceived correctness rather than objective accuracy. However, an important caveat concerns how this asymmetry relates to belief versus desirability. The form of “confirmation” captured here is defined in instrumental terms – i.e. whether feedback confirms one has made the correct choice - rather than whether it aligns with pre-existing beliefs about the world. Unlike other domains (see Tappin et al., 2017), in bandit tasks this distinction is difficult to tease apart because agents typically choose the option they both believe and desire to be better. As a result, the persistence of asymmetric learning from false information could reflect a bias toward belief-consistent evidence, a bias toward desirable outcomes, or some combination of the two. Disentangling these possibilities in the future requires paradigms in which outcome desirability, prior belief, and information reliability are able to be independently manipulated.

An important question raised by these findings concerns the temporal dynamics of the underlying learning process. Participants received two pieces of information sequentially on each trial: first the feedback (win/loss), and then the information accuracy cue (true/false). Given that participants first encountered feedback without knowing whether it was true or false, this raises the question of whether the bias we observe reflects a one or two stage ‘Spinozan’ (Gilbert et al., 1990) process. Under the former, participants would need to maintain feedback in working memory until encountering the accuracy cue and then produce a single integrated update weighted according to both accuracy and whether the information was confirmatory or disconfirmatory. Under the latter, participants would first update in a confirmatory biased manner upon seeing the outcome and then partially revise that update upon seeing the accuracy cue. Crucially, if the initial update is itself biased towards confirmatory evidence, then the residual belief state after a partial revision would inherit this bias, producing the pattern that we observe: attenuated but similarly asymmetric learning from false information.

Whilst an open empirical question, several converging lines of evidence are consistent with a two-stage account. First, confirmation bias has been observed to be present in the absence of any accuracy cues (Palminteri, Lefebvre, et al., 2017), suggesting that asymmetric updating is a default automatic feature of the learning process (Kappes & Sharot 2018) rather than something that requires knowledge of information reliability as a precursor. Second, neuroimaging evidence demonstrates asymmetric coding of positive versus negative prediction errors in the ventral striatum (Lefebvre et al., 2017) at the time of feedback and in the absence of accuracy cues. This suggests that the asymmetry is expressed at the neural level at the time of outcome receipt. Third, a ‘prebunking’ (van der Linden et al., 2017; Roozenbeek & van der Linden, 2019) analogue of the present task - in which participants were told in advance whether upcoming feedback was going to be reliable or not found a similar pattern to us whereby learning was attenuated for unreliable (compared to reliable) feedback but remained significant (Vidal-Perez et al., 2025). The persistence of learning from false information even when falseness is explicitly signalled in advance is consistent with a degree of automaticity in updating at the time feedback is received. Interestingly, these latter results also show evidence of a bias towards confirmatory feedback in the face of unreliable (and reliable) feedback consistent with the results here (with the caveat that the lack of counterfactual feedback in the prebunking design means that this could reflect a positivity or a confirmation bias).

Prior work has shown that asymmetric updating can be normatively advantageous in environments characterised by noise or volatility, for example, by stabilising behaviour or promoting efficient explore/exploit trade-offs (Cazé & van der Meer, 2013; Lefebvre et al., 2022) or when coupled with metacognitive insight (Rollwage & Fleming, 2021), allowing agents to selectively weight information in line with their confidence. More generally, departures from veridical beliefs such as overconfidence (Johnson & Fowler, 2011) and unrealistic optimism (Sharot et al., 2011) have been argued to confer fitness advantages by promoting persistence, decisive action (Johnson & Fowler, 2011) as well as good physical (Rasmussen et al., 2009) and mental (Taylor & Brown, 1988) wellbeing. Seen from this perspective, one possibility is that the persistence of asymmetric learning in the face of explicitly false information may be a by-product of an otherwise adaptive system, one that confers robustness in typical environments but leads to systematic distortions when agents are required to incorporate explicit corrections. Another possibility is that some of the adaptive benefits of asymmetric learning may even persist when outcomes are known to be false (Risen, 2016), analogous to open-label placebo effects that exist even when patients are explicitly informed they are receiving a placebo (Kaptchuk et al., 2010; Locher et al., 2021).

It is worth noting though that the implications of our findings are not confined to the debunking of false information. They extend equally to the verification of true information. Verification and debunking are typically framed as complementary interventions: one confirms accurate claims, the other corrects inaccurate ones. But our results suggest both are subject to the same asymmetry. Just as debunking is least effective when false information is belief-consistent, verification is least effective when true information is belief-inconsistent. Potentially this is because the same mechanism that amplifies learning from confirmatory feedback also attenuates learning from disconfirmatory feedback, regardless of whether the accuracy cue signals true or false. This has an implication for fact-checking efforts: labelling accurate but unwelcome information as verified may do little to shift beliefs precisely in the cases where correction matters the most.

Finally, whilst debates have typically focused on the relative merits of debunking versus prebunking (van der Linden et al., 2017; Roozenbeek & van der Linden, 2019), as well as verifying true versus falsifying false information, our findings highlight a more fundamental constraint: the role of motivational biases in shaping how accuracy labels influence learning. Potentially in real world settings, this asymmetry could be further exacerbated by additional factors such as: identity-relevant beliefs (Kahan et al., 2012; Taber & Lodge, 2006), selective exposure to congruent information (Hart et al., 2009; Stroud, 2010) and limited attention (Kahneman, 2011; Pennycook & Rand, 2019). Together, these findings and our own suggest that the effectiveness of interventions depends not only on the provision of reliable information and cues about accuracy, but also on the broader psychological context, both motivational and attentional, in which that information is processed.

## Data Availability

Data and code available at: https://github.com/HamidRazi7/RL_Misinformation

## Acknowledgements

We thank Nathaniel Daw for the EM code we adapted to fit the models (https://github.com/ndawlab/em/tree/master) to implement the EM algorithm. We also thank Payam Piray for helping with the reparameterization of the model to be able to statistically test for differences in the learning rates.

## Supplemental Materials

### Supplementary Model 1 - Disentangling Confirmation Bias from Positivity Bias

The main text presents Model 4, which operationalises confirmation bias using four learning rates that depend on information accuracy and whether feedback is confirmatory or disconfirmatory. However, the definition of "confirmatory feedback" in Model 4 combines two distinct events: a positive prediction error (PE) for a chosen option (factual outcome) and a negative PE for an unchosen option (counterfactual outcome), and the reverse for disconfirmatory feedback. To check that the pattern or learning does indeed match confirmation bias (Palminteri, Lefebvre, et al., 2017) we ran and expanded version of Model 4 which assigned a unique learning rate to each combination of accuracy, the outcome shown for chosen or unchosen option, and PE sign (negative or positive). This resulted in eight learning rates. The eight learning rates are partitioned according to three trial-by-trial conditions: the Accuracy of the information (cued as True or False), the outcome shown (for the Chosen or Unchosen option), and the sign of the prediction error (Positive or Negative).

For Factual Outcomes (Chosen Option): If Accuracy = True:

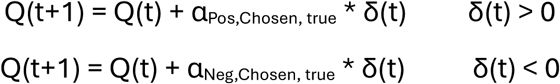

If Accuracy = False:

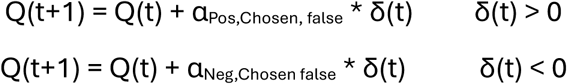

For Counterfactual Outcomes (Chosen Option):

If Accuracy = True:

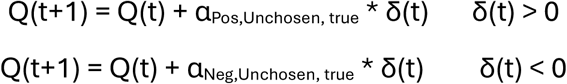

If Accuracy = False:

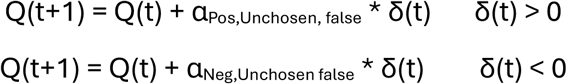

Choice probabilities are generated using the SoftMax function with an inverse temperature parameter β. The model is fitted by minimizing the negative log-likelihood of the participant’s sequence of choices, as described in the main text.

Free parameters (n=9): α_Pos,Chosen, false_, α_Neg,Chosen false,_ α_Pos,Chosen, true_, α_Neg,Chosen, true_, α_Pos,Unchosen, false_, α_Neg,Unchosen false_, α_Pos,Unchosen, true_, α_Neg,Unchosen, true_, β

To test whether learning rates significantly differed between Factual and Counterfactual outcomes, we reparametrized the model such that the unchosen parameters were defined as a subtracted shift (Δ) away from the chosen parameters. For example, in the True trials:

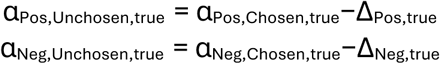

Given that Δ is defined as the difference of α_Chosen_ – α_Unchosen_, by assigning Δ_Pos,true_ and Δ_Neg,true_ as free parameters, the resulting t-statistics for these parameters served as paired tests comparing chosen and unchosen learning rates.

To test the interaction between prediction error sign (positive/negative) and feedback (chosen/unchosen), we reparametrized the model using Average Sums (AS) and Average Differences (AD) of the positive and negative learning rates (detailed derivation is explained later on), where Δ_Interaction_ was a free parameter that tested whether the difference between positive and negative learning rates significantly differed moving from factual to counterfactual outcomes. For example, in the True trials:

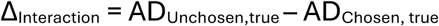

Testing the learning rates for an interaction between prediction error sign (positive/negative) and feedback (chosen/unchosen) separately for true and false feedback revealed a significant interaction in both experiments from true (Experiment 1: t(46) = 5.95, p < 0.001; Experiment 2: t(56) = 3.95, p < 0.001) and from false (Experiment 1: t(46) = 4.68, p < 0.001; Experiment 2: t(56) = 3.92, p < 0.001) feedback.

In Experiment 1 (**Figure 5, Main Text**), this was the result of higher learning rates for positive compared to negative prediction errors when receiving feedback for the option chosen. This was the case both for true feedback (t(46) = 2.59, p = 0.01) and for false feedback (t(46) = 2.20, p=0.03). But we observed the opposite pattern when participants received feedback for the option unchosen, i.e. lower learning rates for positive compared to negative prediction errors. Again, this was the case both for true feedback (t(46) = −2.75, p = 0.008) and for false feedback (t(46) = −2.47, p = 0.01).

In Experiment 2 (Su, the difference between positive versus negative PEs for option chosen feedback was not significant but trending in the right direction both for true (t(56) = 1.88, p = 0.068) and for false (t(56) = 1.86, p=0.067). The difference between positive and negative PEs for the unchosen option feedback was in the opposite direction and trending in the right direction in the case of true (t(56) = −1.87, p = 0.06) and was significant in the case of false (t(56) = −3.81, p < 0.010).

**Supplemental Figure 1:**
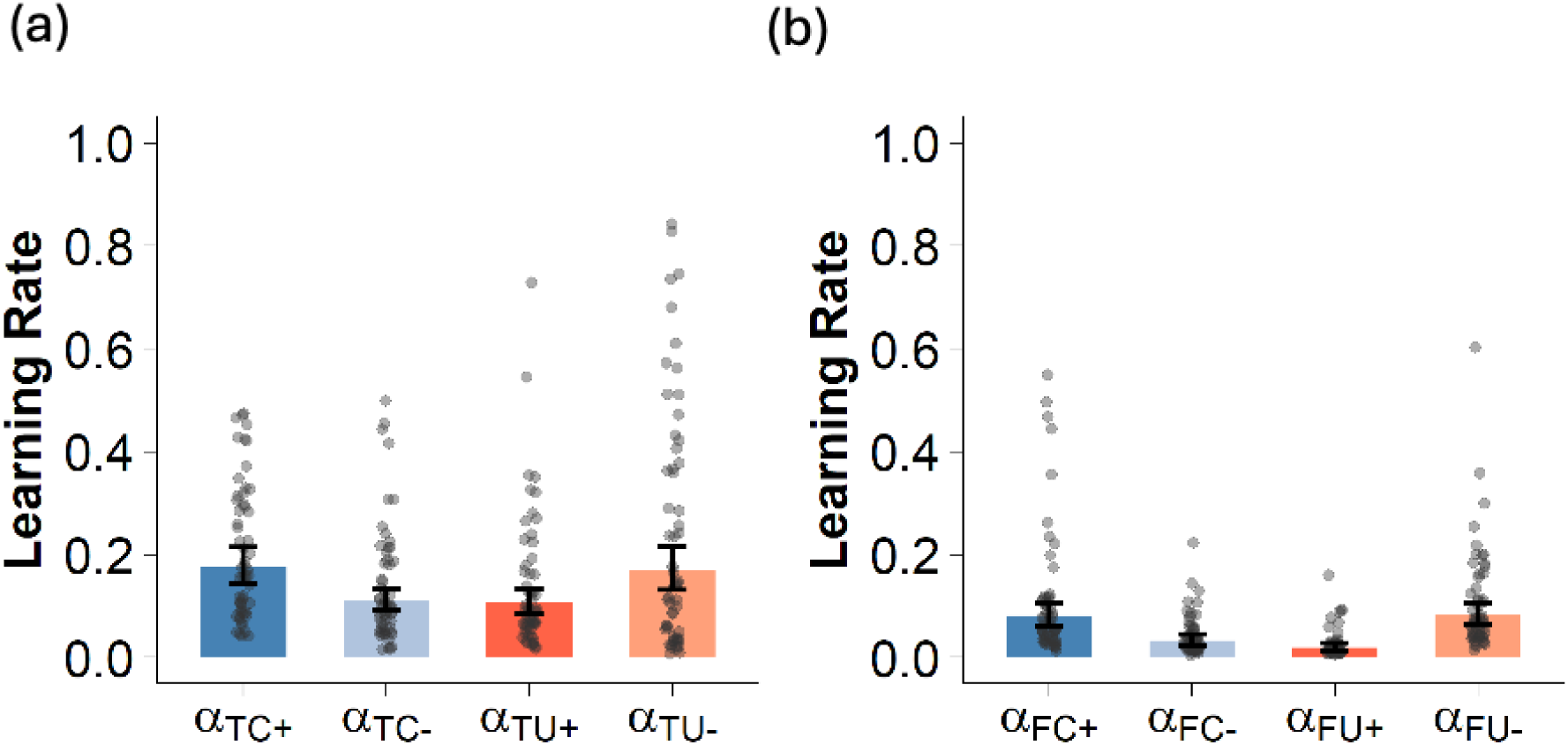
Factual and Counterfactual learning rates from positive and negative prediction errors (Experiment 2). Error bars indicate hierarchical SEM.

### Supplementary Model 2 - Confirmation Bias Across Gain and Loss Contexts

In this model our goal was to see if confirmation bias is robust across Gain (outcomes: +10, +1) and Loss (outcomes: -10, -1) contexts - differences in learning between gains and losses have been reported in the past (Guitart-Masip et al., 2012) but it is unclear whether these impact confirmation bias - so we created separate learning rates for each. Specifically, we modified Model 4 such that the four learning rates either exclusively belonged to the Gain context or to the Loss context.

For the Gain Context:

If Accuracy = True:

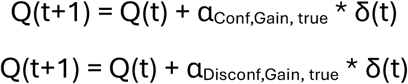

If Accuracy = False:

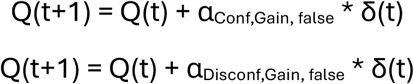

For the Loss Context

If Accuracy = True:

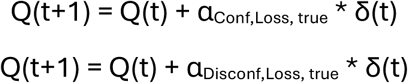

If Accuracy = False:

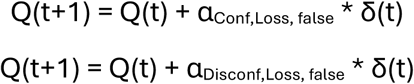

Choice probabilities are generated using the SoftMax function with a single inverse temperature parameter β fitted across both contexts. The model is fitted by minimizing the negative log-likelihood.

Free parameters (n=5) for the Gain version: α_Conf,Gain, false_, α_Disconf,Gain, false,_ α_Conf,Gain, true_, α_Disconf,Gain, true_, β

Free parameters (n=5) for the Loss version: α_Conf,Loss, false_, α_Disconf,Loss, false_, α_Conf,Loss, true_, α_Disconf,Loss, true_, β

**Supplemental Figure 2:**
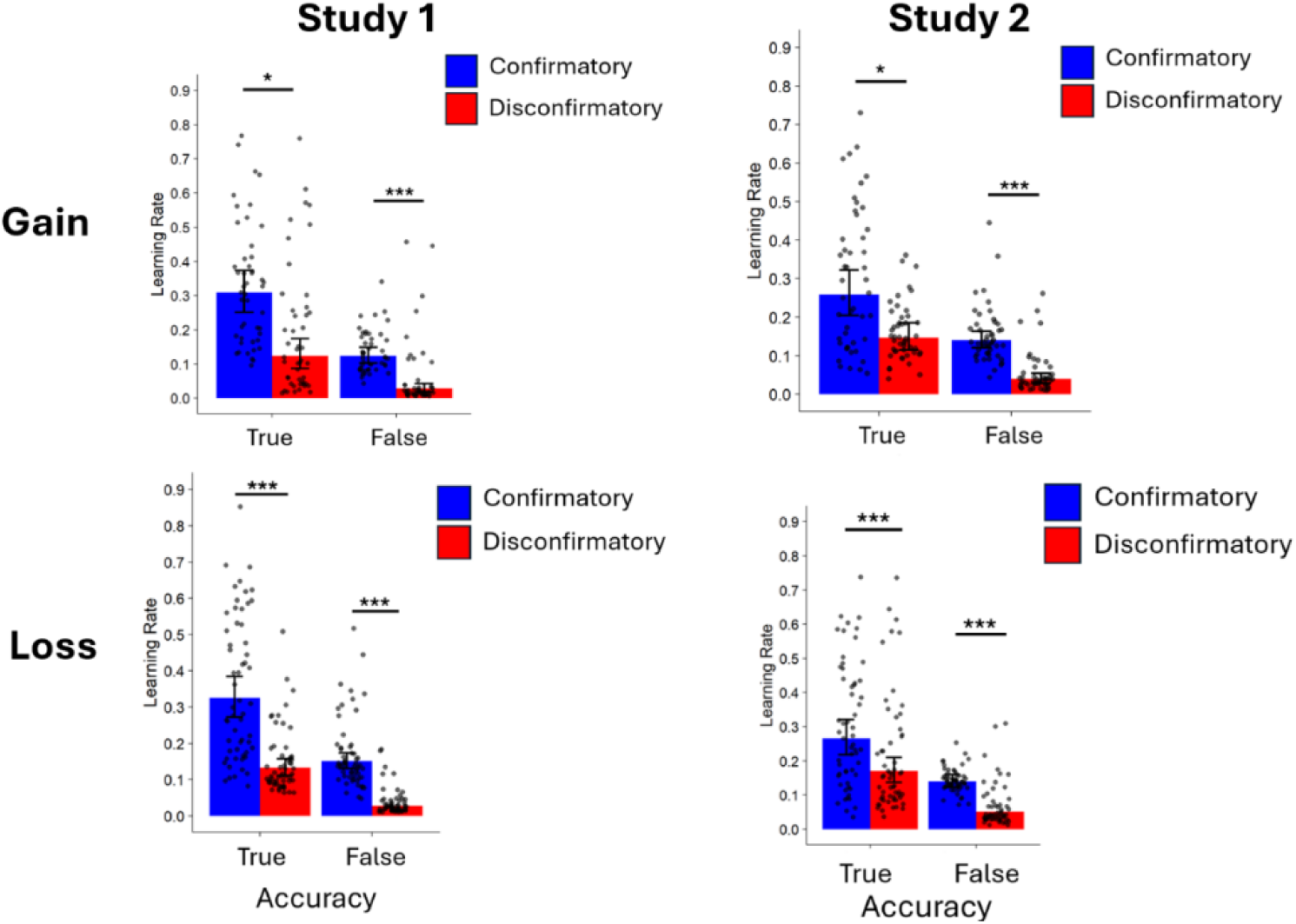
Confirmation bias for Gain and Loss contexts. The bias for true and false information is robust across Gain (above panel) and Loss (lower panel) contexts in both studies. *p < 0.05, ***p < 0.001 hierarchical t-test.

The estimates from this model showed that confirmation bias is robust in the Gain context for both true information (t(46) = 2.51, p = 0.01; t(56) = 2.46, p = 0.01) and false information (t(46) = 3.99, p < 0.001; t(56) = 5.06, p < 0.001). Similarly, the bias exists in the Loss context for both true information (t(46) = 4.05, p < 0.001; t(56) = 3.48, p < 0.001) and false information (t(46) = 4.37, p < 0.001; t(56) = 6.25, p < 0.001).

#### Supplementary Model 3 – Gradual Perseveration

We fit a version of Model 4 which included additional parameters to capture the effects of gradual perseveration (Katahira, 2018; Sugawara & Katahira, 2021; Palminteri, 2023). The core idea behind the gradual perseveration model is to maintain a "choice trace" - like a memory of how often an option has been selected - for both chosen and unchosen options:

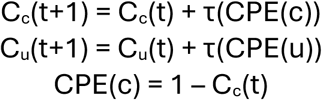

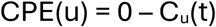

Where C_c_ and C_u_ are the choice traces for the chosen and unchosen options, respectively. When an option is chosen, its trace is increased towards 1; when an option is not chosen, its trace is decreased towards 0. This update process is driven by a “choice prediction error" (CPE) and choice trace accumulation rate (τ) akin to a learning rate, which controls how quickly the trace adapts. For instance, if it is set to 1, then only the previous choice affects the current choice, while lower values mean more of the past choices are influential.

This choice trace then biases future decisions. The predicted probability of picking one option (A) over another (B) on each trial is then determined by a Logit Choice Rule which now includes the choice traces for the respective options (C_A_,C_B_) alongside the Qvalues (Q_A_, Q_B_):

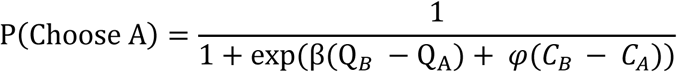

φ (phi) determines how much the history of past choices sways the decision (φ > 0 encourages repeating past choices, i.e. perseverance; φ < 0 encourages switching, i.e. alternation.

The gradual perseveration model therefore has seven free parameters: α_Conf, true_, α_Disconf, true,_ α_Conf, false_, α_Disconf, false_, β, φ, τ.

Comparing the learning rates from this model revealed confirmation bias in the face of false information both in Study 1 (t(46) = 3.13, p < 0.001) and Study 2 (t(56) = 5.24, p < 0.001). Confirmation bias for true information however was only significant in Study 1 (t(46) = 2.43, p = 0.01) but not Study 2 (t(56) = 1.15, p = 0.25).

**Supplemental Figure 3:**
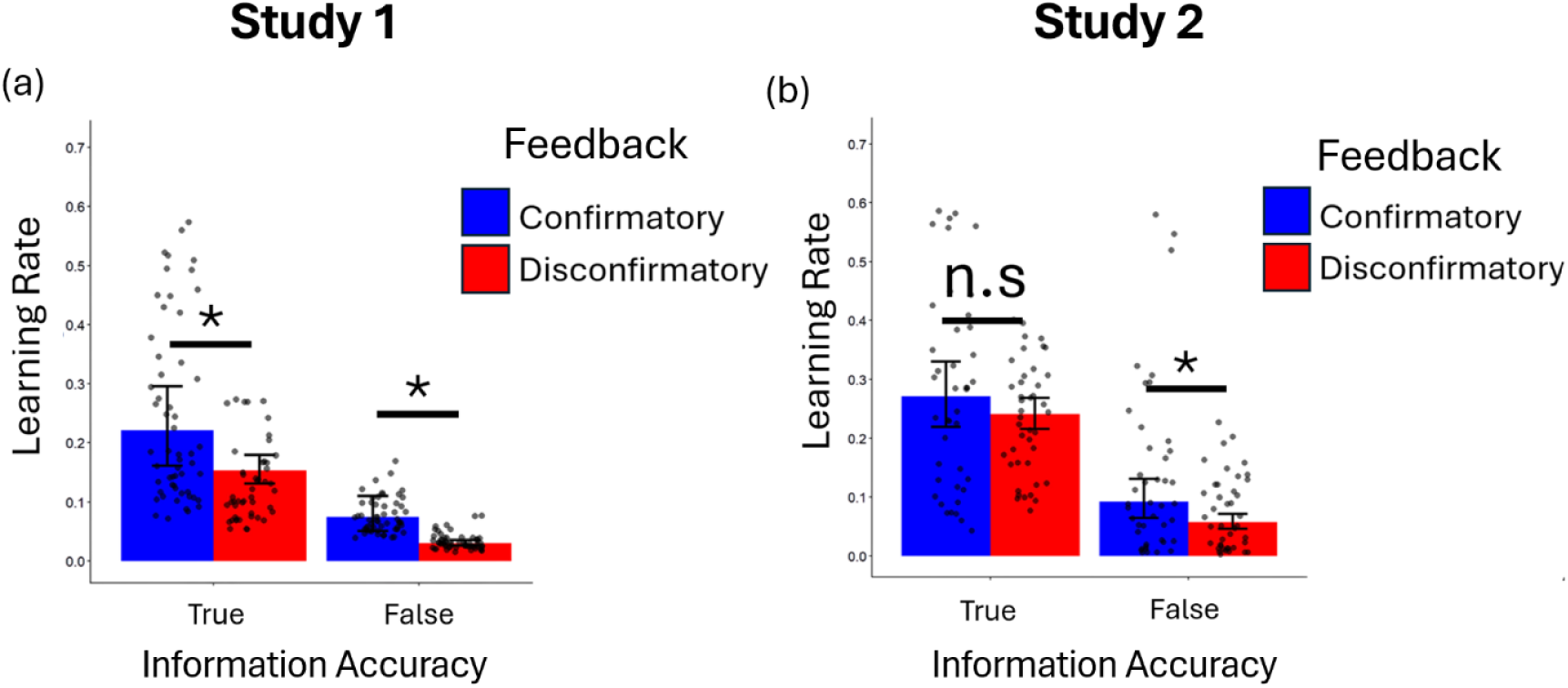
Gradual Perseveration Model Estimates. **(a)** The estimates from a model that includes gradual perseveration again revealed confirmation bias for Study 1 for both true (t(46) = 2.43, p = 0.01) and false (t(46) = 3.13, p < 0.001) information. **(b)** In Study 2, showed confirmation bias remained for false information (t(56) = 5.24, p < 0.001) but not for true (t(56) = 1.15, p = 0.25). n.s: not significant, *p < 0.05, hierarchical t-test. Error bars indicate hierarchical SEM.

### Model Recovery Schematic

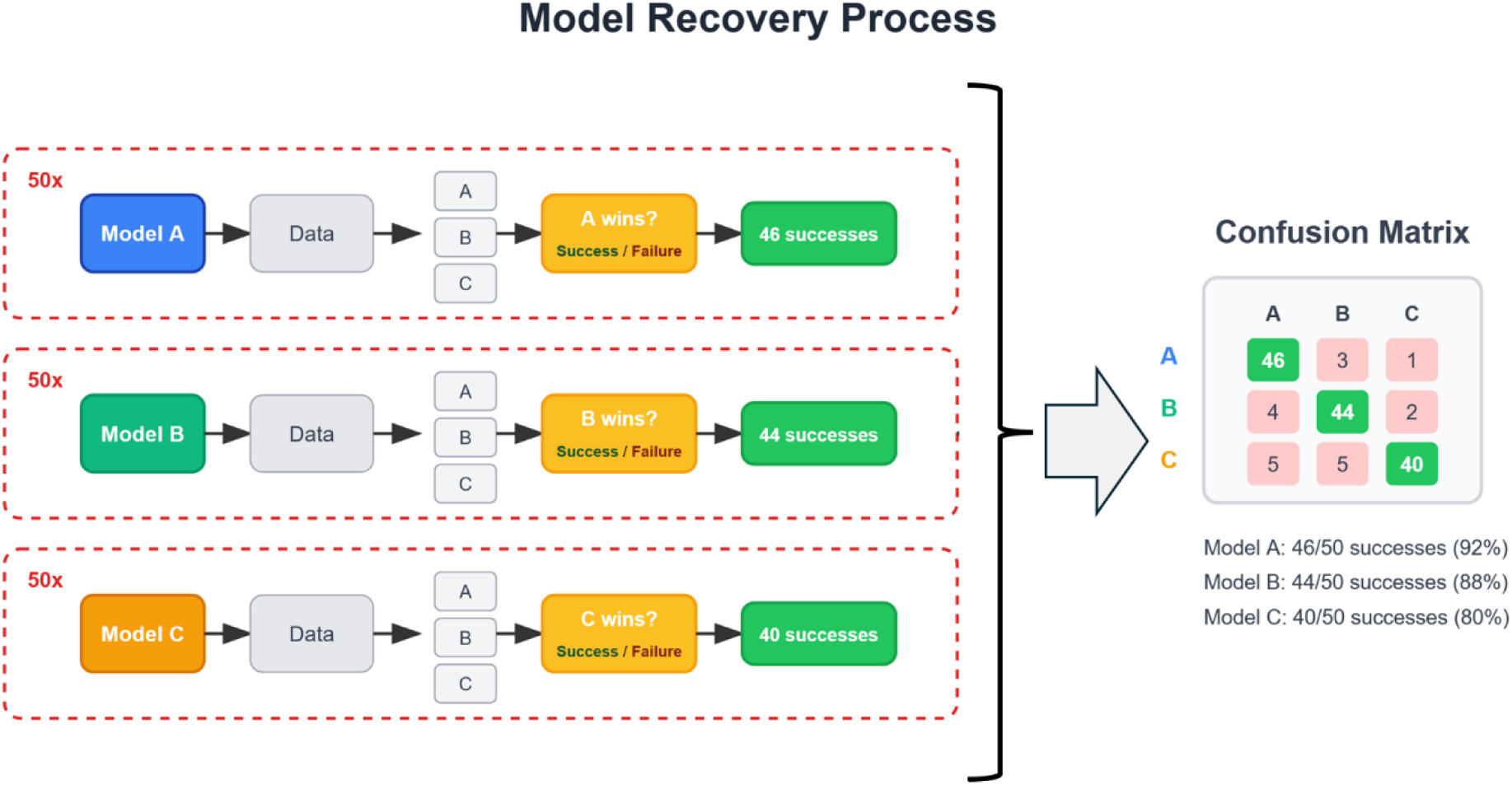

The model recovery process for a hypothetical model space of three models (A, B, and C). The workflow is iterated for all candidate models. In each iteration, one of the models is designated as the ’ground truth’, meaning it is used to simulate the data. For example, the top row shows the process where Model A is the ground truth. A synthetic dataset is generated using Model A and then all models are fitted to this dataset and compared. If Model A comes out as the winner, then this iteration is a success. This entire simulation-and-comparison loop is repeated for 50 iterations, and the number of successes and failures are counted. In the example shown, out of 50 iterations 46 were a success and 4 a failure, corresponding to a 92% recovery rate. The results from all iterations are aggregated into a Confusion Matrix (right panel). In this matrix, the rows represent the true, data-generating model, and the columns represent the model that was selected as the winner by model comparison. The diagonal elements show the proportion of successful recoveries (e.g., the cell A,A shows that Model A was correctly recovered 46 times). The off-diagonal elements show instances of "model confusion," where the model that generated the data did not win (e.g., the cell A,B shows that Model B emerged as the winner 3 times when Model A was the true model). An ideal confusion matrix has high values on the diagonal and low values on the off-diagonals, providing confidence that the models are distinguishable.

### Model Diagnostics

**Supplemental Figure 4:**
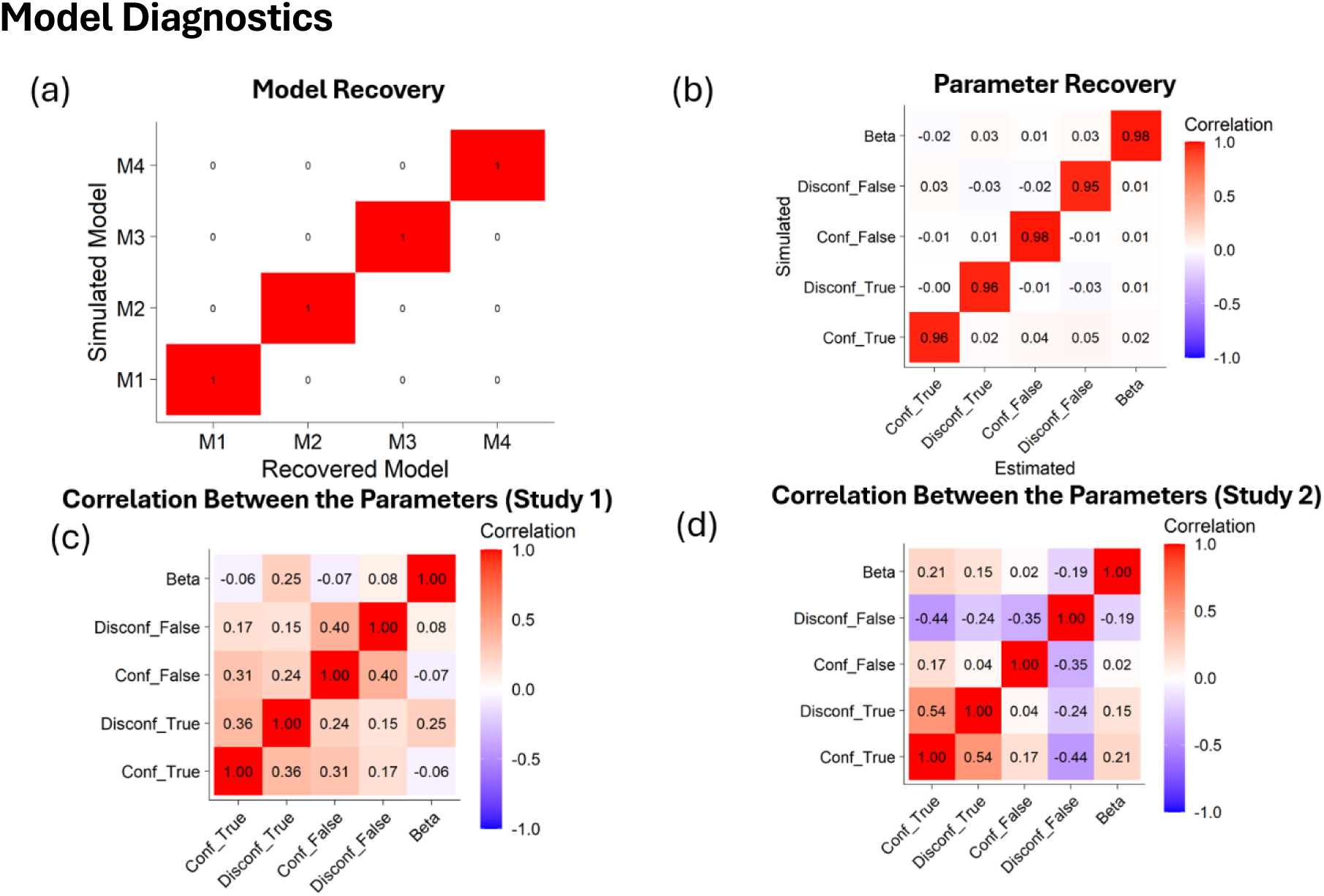
Model Diagnostics. **(a)** Confusion matrix representing model recovery accuracy. All values on the diagonal equal to 1 and off-diagonal values equal to 0, indicating perfect discriminability between the models. **(b)** Heatmap showing Pearson correlation values between parameters used to simulate choice data and parameters estimated from the model fitting process (fit to the same data) for the winning model (M4). **(c)** Heatmap showing the correlation between the actual parameters from the winning model in Study 1 and **(d)** Study 2.

### Parameter Recovery for the Gradual Perseveration Model

**Supplemental Figure 5:**
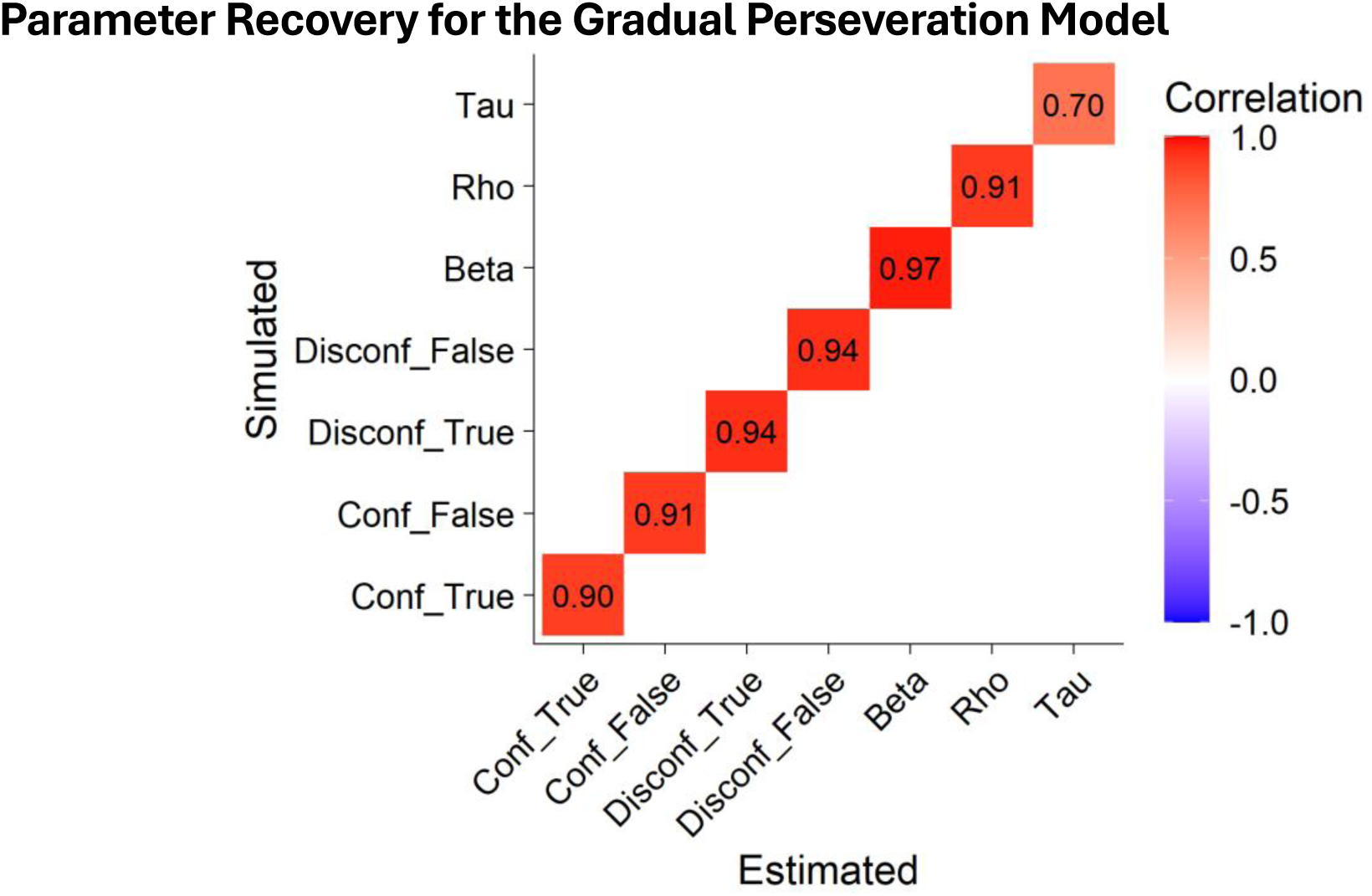
The Gradual Perseveration Model’s Parameter Recovery. The choice trace parameter shows suboptimal recovery, which might make the estimates from this model unreliable.

### Reparameterization to conduct hierarchical t-test and interaction tests

We started reparametrizing Model 4 (M4) by calculating the sum (*AS*_*T*_, *AS*_*F*_) and difference (*AD*_*T*,_ *AD*_*F*_) between confirmatory and disconfirmatory learning rates, separately for true and false trials:

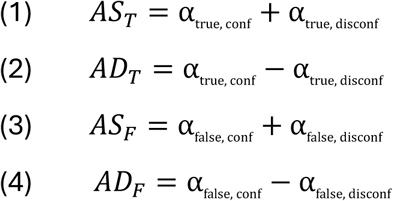

We then added *AS*_*T*_ (from (1)) and *AD*_*T*_ (from (2)) together to get the following equality:

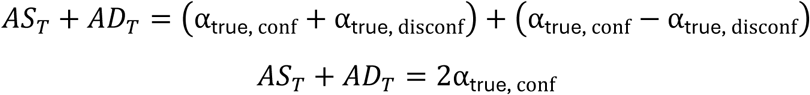

Divided both sides by 2 to get the following expression for α_true, conf_:

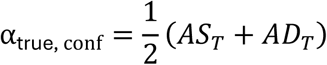

Next, subtracted AD_T_ (from (2)) and AS_T_ (from (1)), then divided by 2 to get a new expression for α_true, disconf_:

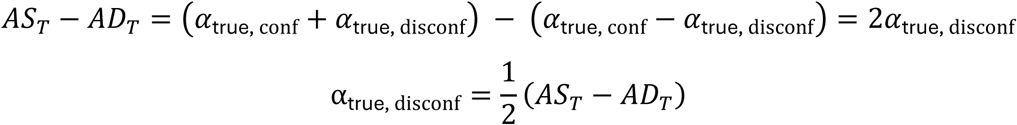

Then applied the same logic to the False learning rates, resulting in the following 4 expressions for the 4 learning rates:

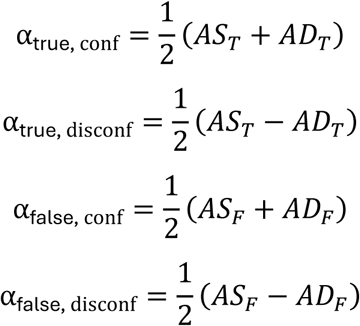

These 4 learning rates are used inside the model, but the free parameters are now: *AS*_*T*_, *AD*_*T*_, *AS*_*F*_, *and AD*_*F*_. Having the model configured in this way then allows us to conduct a hierarchical t-test (against 0) for *AD*_*T*_ which tests the difference between α_true, conf_ and α_true, disconf_ and a hierarchical t-test against 0 for *AD*_*F*_ which tests the difference between α_false, conf_ and α_false, disconf_.

We assessed the reliability of this approach by correlating the estimates from the original model with the estimates from the reparametrized model, which showed a very high correlation for all parameters:

**Supplemental Figure 6:**
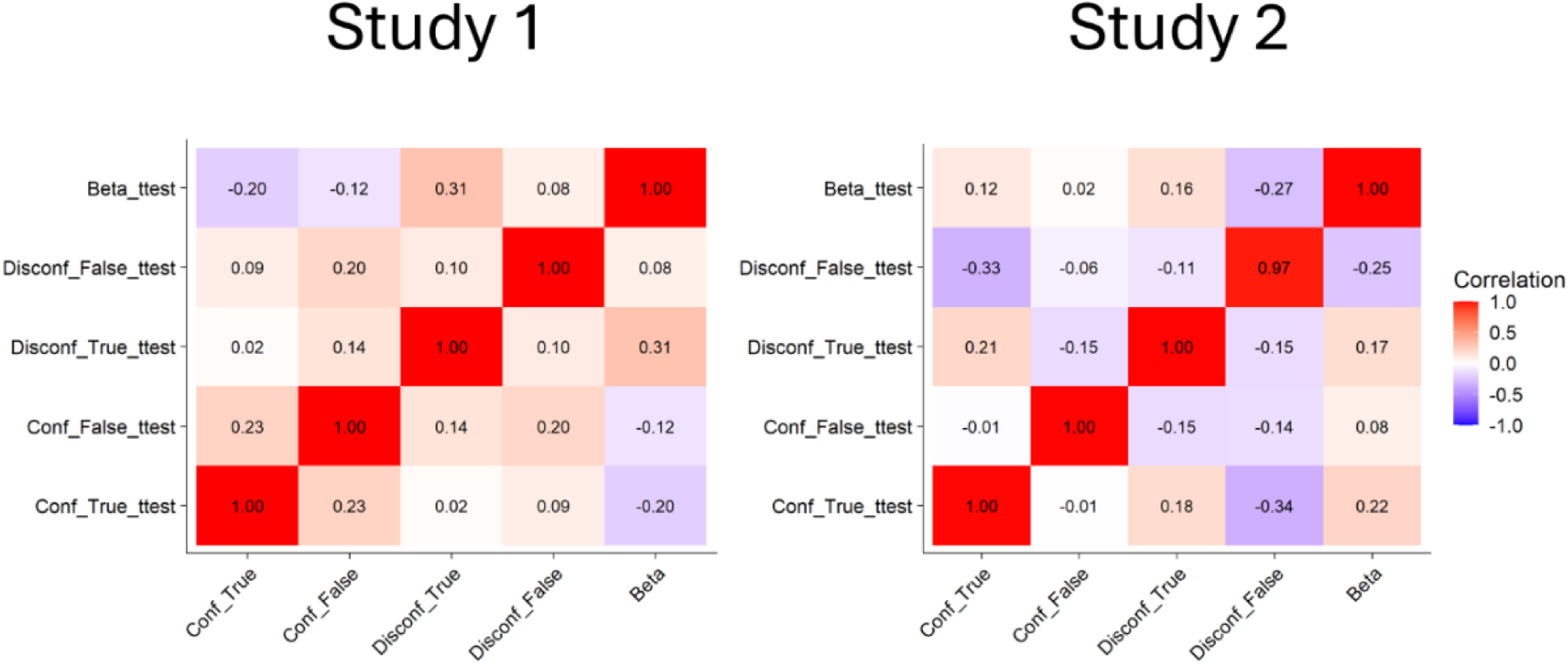
The high reliability of the reparameterization approach is evident in the high correlations between the parameters from the original model and the reparametrized model for the paired t-tests (whose parameters are denoted by _ttest).

Next, we used the same approach and created a reparametrized model for the interaction test. To test whether there is an interaction in learning rates between Feedback (Confirmatory vs Disconfirmatory) and Accuracy (True vs. False), we expressed the interaction as a free parameter, defined as:

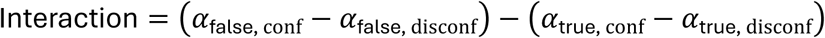

Substitute the expressions:

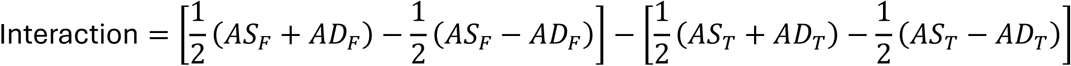

Simplify each term:

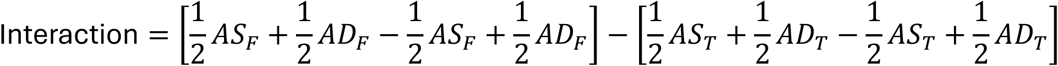

Thus, the interaction simplifies to:

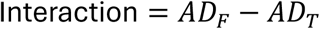

A hierarchical t-test against 0 for Interaction tells us whether there is an interaction between feedback and accuracy or not.

The very high correlation between the reparametrized model and the original model confirmed the reliability of this approach:

**Supplemental Figure 7:**
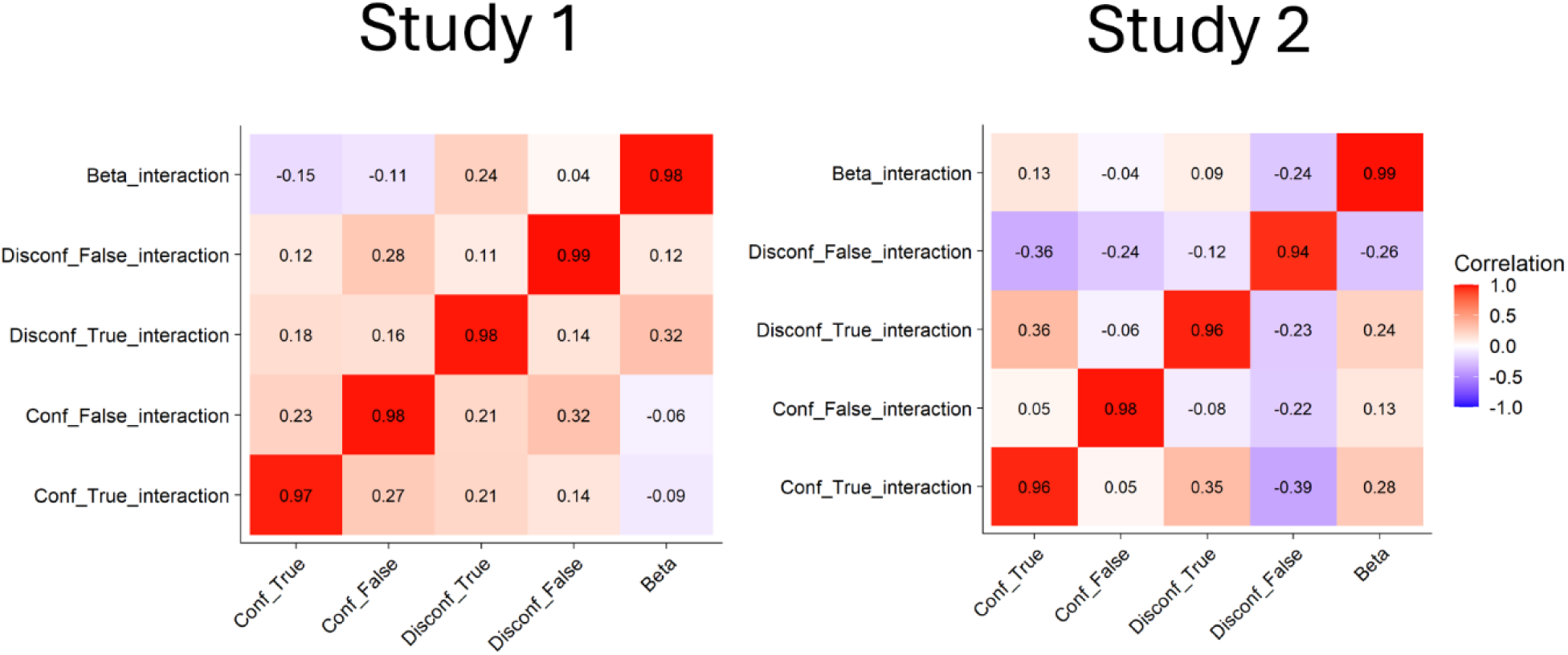
The high reliability of the reparameterization approach is evident in the high correlations between the parameters from the original model and the reparametrized model for testing the interaction (whose parameters are denoted by _interaction).

